# Multi-night EEG reveals positive association between sleep efficiency and hippocampal subfield volumes in healthy aging

**DOI:** 10.1101/2023.11.05.565729

**Authors:** Attila Keresztes, Éva Bankó, Noémi Báthori, Vivien Tomacsek, Virág Anna Varga, Ádám Nárai, Zsuzsanna Nemecz, Ádám Dénes, Viktor Gál, Petra Hermann, Péter Simor, Zoltán Vidnyánszky

**Author notes:** Corresponding Author: Attila Keresztes, Research Centre for Natural Sciences, Magyar, tudósok körútja 2, 1117, Budapest, Hungary. These authors contributed equally to this publication (Attila Keresztes, Éva Bankó).

## Abstract

Age-related atrophy of the human hippocampus and the enthorinal cortex starts accelerating at around age 60. Due to the contributions of these regions to many cognitive functions seamlessly used in everyday life, this can heavily impact the lives of elderly people. The hippocampus is not a unitary structure and mechanisms of its age-related decline appear to differentially affect its subfields. Human and animal studies have suggested that altered sleep is associated with hippocampal atrophy. Yet, we know little about subfield specific effects of altered sleep in healthy aging and their effect on cognition. Here, in a sample of 118 older adults (M_age_ = 63.25 years), we examined the association between highly reliable hippocampal subfield volumetry, sleep measures derived from multi-night recordings of portable electroencephalography and episodic memory. Objective sleep efficiency – but not self-report measures of sleep – was associated with entorhinal cortex volume when controlling for age. Age-related differences in subfield volumes were associated with objective sleep efficiency, but not with self-report measures of sleep. Moreover, older adults characterized by a common multivariate pattern of subfield volumes that contributed to positive sleep– subfield volume associations, showed lower rates of forgetting. Our results showcase the benefit of objective sleep measures in identifying potential contributors of age-related differences in brain-behavior couplings.

## main text

Volume loss of the hippocampus and surrounding mediotemporal cortical areas during human aging is accelerated relative to most other cortical areas (Fjell et al., 2009; Raz et al., 2010; Pomponio et al., 2020). In vivo animal and post-mortem human studies have suggested that these volumetric changes can reflect a combination of factors (for a comprehensive review, see Bettio et al., 2017) including decreased number, size and density of axons and dendrites due to synaptic pruning, to some extent decreased number of neurons (West, 1993; Lister and Barnes, 2009), as well as changes in astrocytic and glial cell size and density (Ojo et al., 2015). These alterations, in turn, are thought to be the consequences of genetically (Guo et al., 2019) and environmentally (Binnewies et al., 2023) driven changes in blood supply (Shing et al., 2011), in hormonal (McEwen, 1997), neurotrophic (Buhusi et al., 2017), and neuron-glia interaction regulation (Ojo et al., 2015), and in neurogenesis (Spalding et al., 2013; Boldrini et al., 2018; cf. Sorrells et al., 2018).

Understanding the mechanisms of these changes is crucial for mitigating or potentially reversing (McEwen, 1997; Kim et al., 2016; Bettio et al., 2017) their adverse effects on cognitive functions supported by the hippocampus. Because these functions – including memory (Wixted and Squire, 2011; Moscovitch et al., 2016), navigation (Eichenbaum, 2017; Whittington et al., 2020), and even language (Duff et al., 2020) – have major roles in everyday life, their impairment has a significant impact in healthy, and devastating effect in pathological aging. Critically, hippocampal volumetry is currently the only widely accessible, valid, and reliable method to assess hippocampal atrophy in humans *in vivo*.

Importantly, age-related changes driving volumetric loss of the hippocampus appear to be expressed differentially across its subfields (Bartsch and Wulff, 2015) varying in cytoarchitecture, macrostructure, connectivity (Amaral and Lavenex, 2007), and function (Mueller et al., 2011; Genon et al., 2021). This heterogeneity may explain the fact that although hippocampal volume loss has been consistently detected in aging, no consistent trajectories of hippocampal subfield volume loss have been identified (de Flores et al., 2015). Thus, for a mechanistic understanding of hippocampal atrophy in healthy aging it is crucial to tease apart subfield specific contributions of factors affecting hippocampal volume.

One such relatively understudied factor is sleep. It is well documented that sleep patterns altered in apnea (Macey et al., 2002; Kim et al., 2016), insomnia (Noh et al., 2012), narcolepsy (Joo et al., 2012), as well as in pathological conditions such as depression (Campbell et al., 2004; Videbech and Ravnkilde, 2004) are associated with smaller total hippocampal volume. Recent studies have also provided evidence suggesting that sleep disturbances may partly underlie the effects of pathological (Elcombe et al., 2015; Burke et al., 2022) and healthy (Carvalho et al., 2017; Liu et al., 2018; Alperin et al., 2019; Fjell et al., 2020) aging on hippocampal volume with subfield specific effects (Joo et al., 2014; Lam et al., 2021; Liu et al., 2021; De Looze et al., 2022). The absence of observed sleep-volume associations when only total hippocampal volumes are assessed underscores the need for subfield specific studies (Sabeti et al., 2018; De Looze et al., 2022).

To date, only two studies have assessed hippocampal subfield volumes as a function of sleep in healthy aging. In a large sample (n = 417) De Looze and colleagues (2022) found that both too short and too long sleep duration were associated with smaller subiculum volume, with too long sleep additionally associated with smaller volumes of the cornu ammonis 1 (CA1) subfield. Somewhat consistently, Liu and colleagues (Liu et al., 2021) in a smaller sample study (n = 67) found that healthy participants with poor sleep had smaller volumes of subiculum, CA1, and dentate gyrus. Importantly, both studies relied on self-report measures to assess sleep quality and duration, thus it remains to be clarified to what extent the observed associations with hippocampal subfield volumetry reflect alterations of objective sleep parameters. Furthermore, only the Liu et al. (2021) study used submillimeter resolution magnetic resonance imaging (MRI) to perform hippocampal segmentations, with various arguments supporting the limited validity of lower resolution-based segmentations (Wisse et al., 2014, 2021). Finally, no studies have yet investigated the potential effect of sleep-related age-differences in hippocampal subfields on memory.

To address these gaps we examined the association between highly reliable semi-automatic hippocampal subfield volumetry and sleep measures derived from multi-night recordings of portable electroencephalography (EEG) and their association with standard measures of learning and memory in healthy aging.

Participants were recruited as part of the Hungarian Longitudinal Study of Healthy Brain Aging (HuBA) at the Brain Imaging Centre of the Research Centre for Natural Sciences in Budapest, Hungary. They had no previous or current diagnosis of neurological or psychiatric disorders, untreated hypertension, diabetes and no history of alcohol or drug abuse or malignant tumor in the past 5 years. Data of one hundred and eighteen 50–80-year-old participants (M_age_ = 63.25, SD_age_ = 7.23, 68 females, M_education_ = 16.25, SD_education_ = 2.55) who had hippocampal subfield data (n=112) or sleep recordings for at least 3 nights (n=61), was included in this study. Of these, 55 participants (M_age_ = 62.64, SD_age_ = 6.85, 30 females) had both hippocampal subfield and sleep data. The study was approved by the National Institute of Pharmacy and Nutrition (OGYÉI/68903/2020), and all participants gave written informed consent.

Using methods described in detail previously (Keresztes et al., 2020, 2022), we acquired high-resolution (0.4 mm × 0.4 mm × 2.0 mm) MRI images of the hippocampus and surrounding mediotemporal areas perpendicular to the longitudinal axis of the right hippocampus on a 3 T Siemens Magnetom Prisma scanner. Then, in a semi-automatic procedure, we used the Automated Segmentation of Hippocampal Subfields (ASHS) (Yushkevich et al., 2015) software package to delineate four regions of interest bilaterally based on a custom lifespan atlas (Bender et al., 2018): three within the hippocampal body – subiculum (SUB), CA regions 1 and 2 (CA1-2), and dentate gyrus–CA3 (DG-CA3) – as well as the entorhinal cortex (EC) on six consecutive slices anterior to the hippocampal body (see Figure 1A). Hippocampal body ranges were manually defined by a rater (V.A.V.) with good inter-(all Cohen’s kappas > 7) and intra-rater reliability (all Cohen’s kappas > .77). Segmentations were visually inspected by both raters who then corrected obvious errors. Volumes for all regions were adjusted for intracranial volume, estimated by ASHS, based on the analysis of covariance approach (Jack et al., 1989; Raz et al., 2005), then summed across hemispheres for all analyses.

**Figure 1.**
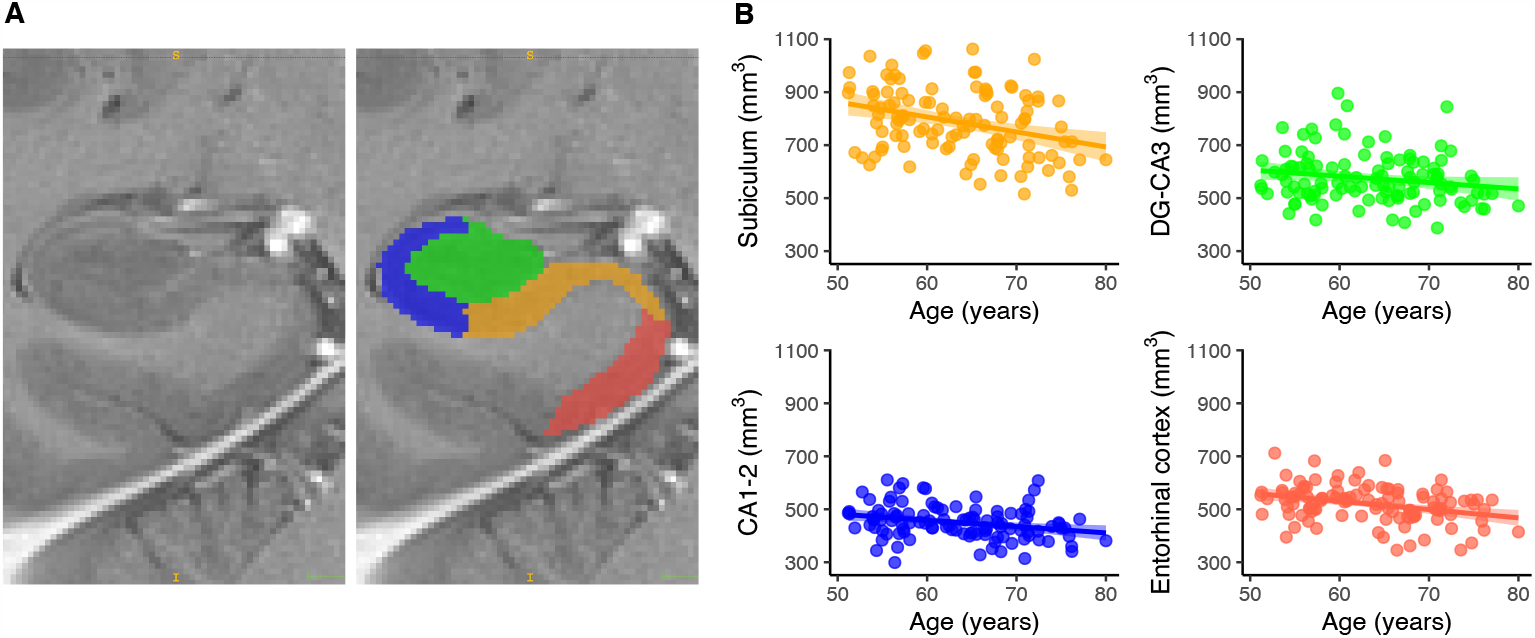
(A) Example hippocampal subfield segmentation of the right hippocampus and entorhinal cortex into four subfields: subiculum (orange), dentate gyrus – Cornu Ammoni 3 (DG-CA3; green), CA1-2 (blue), and entorhinal cortex (red). (B) Negative age-associations of the four subfields, non significant only for DG-CA3.

Sleep recordings were acquired at participants’ homes using a wireless Dreem2 EEG headband (Dreem, Rythm, Paris, France; DH) with five electrode sensors yielding seven bipolar derivations at a 250 Hz sampling frequency. Participants wore the DH for at least 7 consecutive nights starting an average of 29.9 days ([2–193, SD = 33.6] after the MRI session. Sleep recordings were qualified and scored by a deep learning algorithm which previously demonstrated equivalent performance to human raters (Arnal et al., 2020). Data above 74.5 % quality index for at least one bipolar derivation was retained. Recordings with unusual length or rapid eye movement (REM) sleep percentage above 40% were additionally reviewed for inclusion by a sleep expert. This procedure yielded an average of 5.98 nights (*SD* = 1.47, *Mdn* = 7).

We used the proportion of the time spent asleep and the time spent in bed (objective sleep efficiency) as an independent variable. To assess subjective estimates of sleep efficiency and quality, we used three measures derived from self-report questionnaires. First, we calculated subjective sleep efficiency similarly to the objective sleep efficiency using items 1,3, and 4 of the Pittsburgh Sleep Quality Index (PSQI; Buysse et al., 1989; Hungarian adaptation: Takács et al., 2016). Second, we assessed subjective sleep quality for the past month using the Athens Insomnia Scale (AIS; Soldatos et al., 2000; Hungarian adaptation: Novák, 2004). The 8-item AIS assesses the severity of sleep problems based on the insomnia criteria defined by the International Classification of Diseases (ICSD-10; ICSD: American Academy of Sleep Medicine, 2005) with a total score of 6 or above indicating the presence of insomnia symptoms. We used the 14-item Groningen Sleep Quality Scale (GSQS; Meijman et al., 1988; Hungarian adaptation: Simor et al., 2009) to assess the subjective quality of sleep each morning, after awakening. Higher scores in the GSQS reflect higher subjective sleep fragmentation. We modified the original binary (yes/no) scoring of the GSQS to a 5-point Likert scale to weight each statement and used an average of GSQS scores across all nights. All reported *p* values are non-corrected for multiple comparisons.

Bivariate zero-order Pearson’s correlations revealed significant negative age-volume associations (Figure 1B) for SUB (*r*_*(110)*_ = -.32, *p* < .001), CA1-2 (*r*_*(110)*_ = -.27, *p* = .004), and EC (*r*_*(110)*_ = -.34, *p* < .001), but not DG-CA3 (*r*_*(110)*_ = -.18, *p* = .055), as well as significant negative age-sleep associations for objective sleep efficiency (*r*_*(53)*_ = -.39, *p* = .002), but not for subjective measures of sleep quality or duration (all *r*s < .17, all *p*s > .139).

We assessed the association between objective sleep efficiency and hippocampal subfield volumes in regression models defined for each subfield volume controlling for age, sex and education. In addition, each model included covariates for physical activity (total metabolic equivalent minutes per week measured by the short version of the International Physical Activity Questionnaire (Craig et al., 2003)), total IQ (measured by the Wechsler Adult Intelligence Scale 4^th^ Edition (WAIS-IV; Pearson PLC, London, UK)), depression (measured by the short version of the Geriatric Depression Scale (GDS;Sheikh and Yesavage, 1986), and blood pressure (systolic and diastolic blood pressure measured after each scanning session). Only the model predicting entorhinal cortex volume provided a significant fit to the data (*F*(9,37) = 4.54, *p* < .001, *R*^*2*^ = .41) with objective sleep efficiency (*β* = 7.18, *p* = .004) and age (*β* = -4.04, *p* = .008) significantly predicting entorhinal cortex volume (Figure 2). Models for all other hippocampal subfield volumes were non-significant (all *F*s < 1.18, and all *p*s > .1).

**Figure 2.**
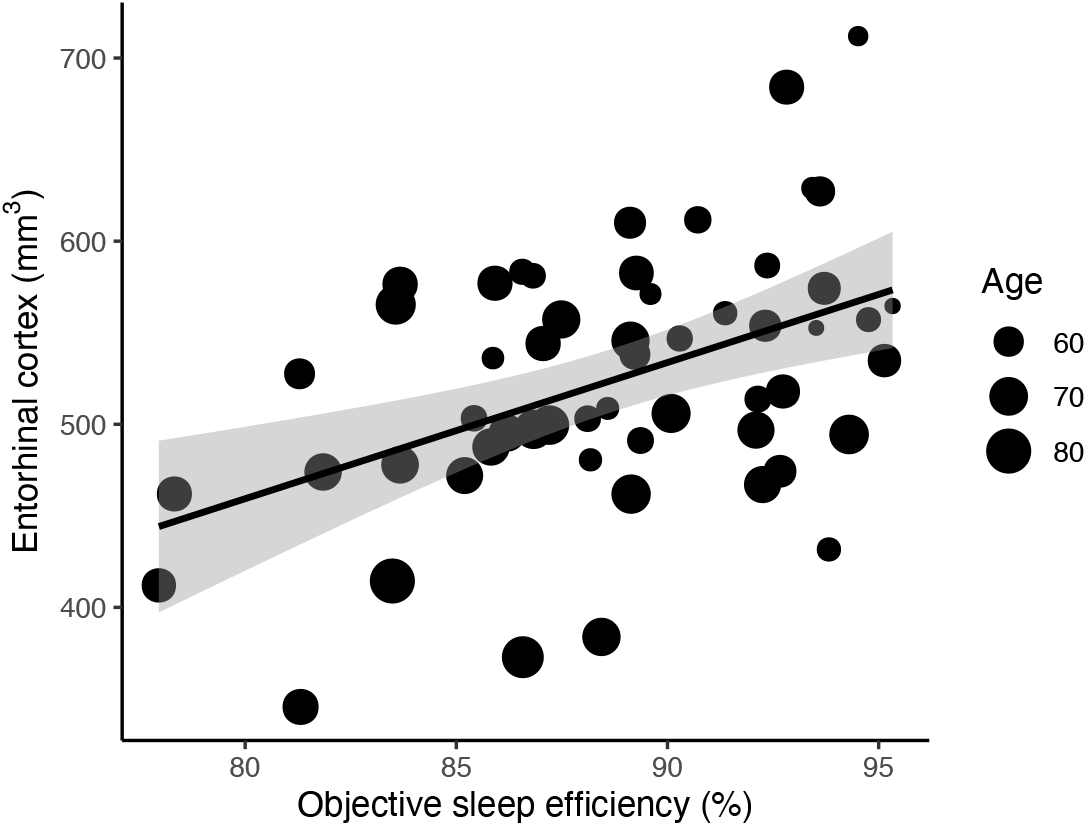
Entorhinal cortical volume positively associated with objective sleep efficiency.

We used identical models to test associations between self-report measures of sleep and hippocampal subfields. For all three measures only the model predicting entorhinal cortex was significant and only for the GSQS: *F*(9,43) = 2.46, *p* = .023, *R*^*2*^ = .2 and the AIS: *F*(9,53) = 3.01, *p* = .006, *R*^*2*^ = .23 with only age significantly predicting volume (GSQS: *β* = -4.97, *p* = .001, AIS: *β* = -5.27, *p* < .001).

Next, we assessed whether the combined pattern of age-related differences in hippocampal subfield and entorhinal cortex volumes were associated with any sleep measure. To this end, we used partial least squares correlations (PLSC) (Krishnan et al., 2011), a robust method that allows detecting multivariate associations between groups of variables (McIntosh and Lobaugh, 2004). Following methods described in detail in Keresztes et al., (2017), we used singular value decomposition to calculate a single latent variable (LV) – significant as assessed by permutation tests with 10000 iterations, *p* < .001 that expresses the largest common variance between age and volumetric measures, with the significance of each volume’s contribution to the LV (weight) assessed by 10000 bootstrapped resampling of the data. This analysis revealed that all volumetric measures contributed significantly (all *p*s <.029) to the LV with entorhinal cortex and subiculum (for both z > 3.9, *p* < .001) driving the LV–age association most strongly (Figure 3A). By multiplying the weight vector of volumetric measures with each participant’s vector of observed volumes, we obtained a single score for each participant that reflects how well that participant’s data fits the common pattern of observed age-related differences in hippocampal subfield volumes. Pearson’s correlations revealed that this score was significantly negatively associated (*r*_*(53)*_= -.42, *p* =.001) with objective sleep efficiency (Figure 3B), but showed no associations with any of the subjective sleep measures (all *r*s < .16, all *p*s > .25).

**Figure 3.**
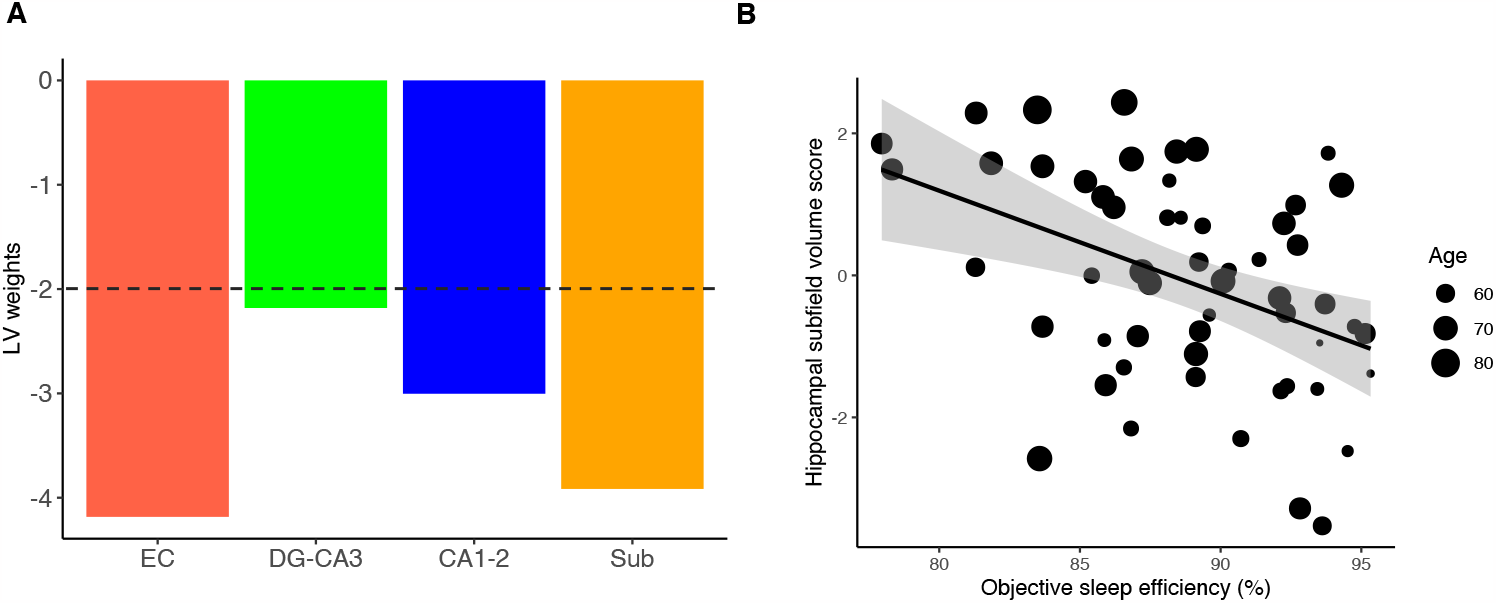
(A) Subfields specific weights of the latent variable (LV) expressing the largest common variance between age and volumes of the four subfields. Bootstrapped values can be interpreted as z-values, with values >2 or <-2 (represented by the dashed line) considered significant. EC: entorhinal cortex, Sub: subiculum. (B) A hippocampal subfield volume score for each participants reflects how much that individual’s data represent the LV, i.e., the multivariate subfield volume – age association. This score is negatively associated with objective sleep efficiency.

Finally, we assessed whether the associations between sleep and hippocampal subfield volumes manifest in individual differences in memory. To this end, we used PLSC to (i) identify potential LVs that express common patterns of variance between age-residualized measures of hippocampal subfield volumes and sleep quality, and to (ii) test the association between individual differences in the expression of these LVs and in delayed recall on the Rey Auditory Verbal Learning Test (RAVLT; Lezak, 2004; Hungarian adaptation: Kónya et al., 1995). This analysis (see Figure 4) identified one significant LV (*p* = .0174), expressing a positive association between sleep and hippocampal subfield volume, with entorhinal cortex (*p* < .001) and subiculum (*p* = .042) contributing significantly (Figure 4A) to positive associations with objective sleep efficiency and AIS (Figure 4B). Again, we also calculated a single score for each participant that reflects how well that participant’s data fit the common pattern of observed differences in hippocampal subfield volumes that contribute to a positive association with sleep measures (independently of age). Importantly, this score correlated positively at a trend level (*r*_*(51)*_= .27, *p* = .05) with age-residualized measures of delayed recall (percent of learnt items recalled at a 30-min delay) and significantly negatively with forgetting (number of forgotten items after a 30-min delay), *r*_*(51)*_= -.31, *p* = .023 (Figure 4C).

**Figure 4.**
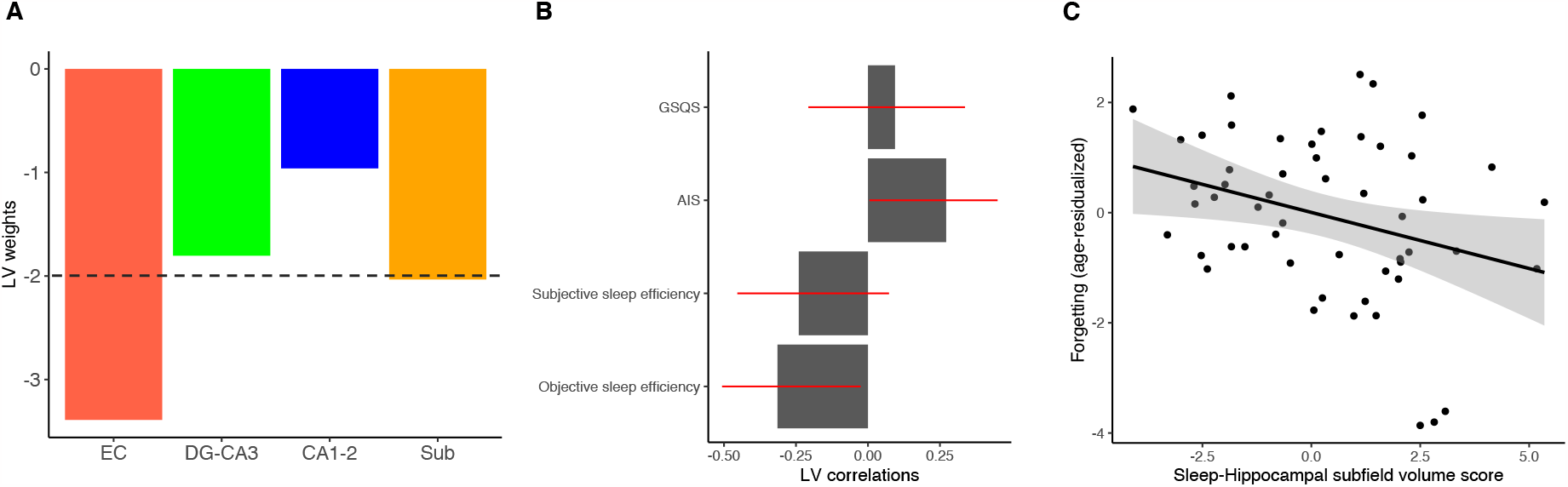
(A) Subfields specific weights of the latent variable (LV) expressing the largest common variance between age-residualized volumetric and sleep measures. Bootstrapped values can be interpreted as z-values, with values >2 or <-2 (represented by the dashed line) considered significant. EC: entorhinal cortex, Sub: subiculum. (B) Correlations of sleep measures with the LV in the optimal solution of the PLSC. Red lines depict 95% confidence intervals (C) The Sleep-Hippocampal subfield volume score for each participants reflects how much that individual’s data represent the LV, i.e., the multivariate subfield – sleep association. This score is negatively associated with forgetting after a 30-min delay.

In sum, multi-night sleep EEG and high-resolution structural MRI of the mediotemporal lobe revealed that objective sleep efficiency was associated with volumes of the entorhinal cortex in a sample of healthy older adults when controlling for age. Importantly, age-related differences in subfield volumes – mainly driven by entorhinal cortical and subiculum volume – were associated with objective sleep efficiency, but not with self-report measures of sleep. Moreover, at the individual level, older adults characterized by a common multivariate pattern of subfield volumes that contributed to positive hippocampal subfield volume – sleep associations, performed better on a standard delayed recall.

Together, these results provide converging evidence for the notion that sleep is associated with age-related changes in brain–behavior couplings. It also showcases the importance of considering objective sleep parameters for studies of these associations. Critically, many of the reported sleep associations were detected via sleep EEG recorded over multiple nights but not with self-report measures. Thus, our results also highlight the sensitivity of an easy-to-use tool with high ecological validity that could provide objective measures of sleep quality.

Some limitations of our study are worth noting. First, due to concerns of validity (Wisse et al., 2017), and in line with our prior work across the lifespan (Bender et al., 2018; Keresztes et al., 2020), we only assessed hippocampal subfield volumes along the hippocampal body, and assessed entorhinal cortical volume on six slices anterior to it. Second, compared to gold standard polysomnography, DH recordings, albeit easy-to-use and require less resources, provide less reliable data, both in terms of accuracy and data loss due to technical difficulties. Third, the cross-sectional age-associations presented here cannot allow inferences about change and sleep–brain couplings over time (Raz and Lindenberger, 2011; cf., Keresztes et al., 2022).

To conclude, the study presented showcases that objective sleep measures can reveal otherwise unnoticed associations that are potentially important contributors of age-related differences in brain-behavior couplings. Future studies are needed to reveal subfield specific mechanisms driving these associations. In addition, our results call for longitudinal studies to allow for inferences about actual change.

